# Development of a soft cell confiner to decipher the impact of mechanical stimuli on cells

**DOI:** 10.1101/2020.01.04.892695

**Authors:** A. Prunet, S. Lefort, H. Delanoë-Ayari, B. Laperrousaz, G. Simon, S. Saci, F. Argoul, B. Guyot, J.-P. Rieu, S. Gobert, V. Maguer-Satta, C. Rivière

## Abstract

Emerging evidence suggests the importance of mechanical stimuli in normal and pathological situations for the control of many critical cellular functions. While the effect of matrix stiffness has been and is still extensively studied, few studies have focused on the role of mechanical stresses. The main limitation of such analyses is the lack of standard *in vitro* assays enabling extended mechanical stimulation compatible with dynamic biological and biophysical cell characterization. We have developed an agarose-based microsystem, the soft cell confiner, which enables the precise control of confinement for single or mixed cell populations. The rigidity of the confiner matches physiological conditions and enables passive medium renewal. It is compatible with time-lapse microscopy, *in situ* immunostaining, and standard molecular analyses, and can be used with both adherent and non-adherent cell lines. Cell proliferation of various cell lines (hematopoietic cells, MCF10A epithelial breast cells and HS27A stromal cells) was followed for several days up to confluence using video-microscopy and further documented by Western blot and immunostaining. Interestingly, even though the nuclear projected area was much larger upon confinement, with many highly deformed nuclei (non-circular shape), cell viability, assessed by live and dead cell staining, was unaffected for up to 8 days in the confiner. However, there was a decrease in cell proliferation upon confinement for all tested cell lines. The soft cell confiner is thus a valuable tool to decipher the effect of long-term confinement and deformation on the biology of cell populations. This tool will be instrumental in deciphering the impact of nuclear and cytoskeletal mechanosensitivity in normal and pathological conditions involving highly confined situations, such as those reported upon aging with fibrosis or during cancer.

**Graphical abstract:** 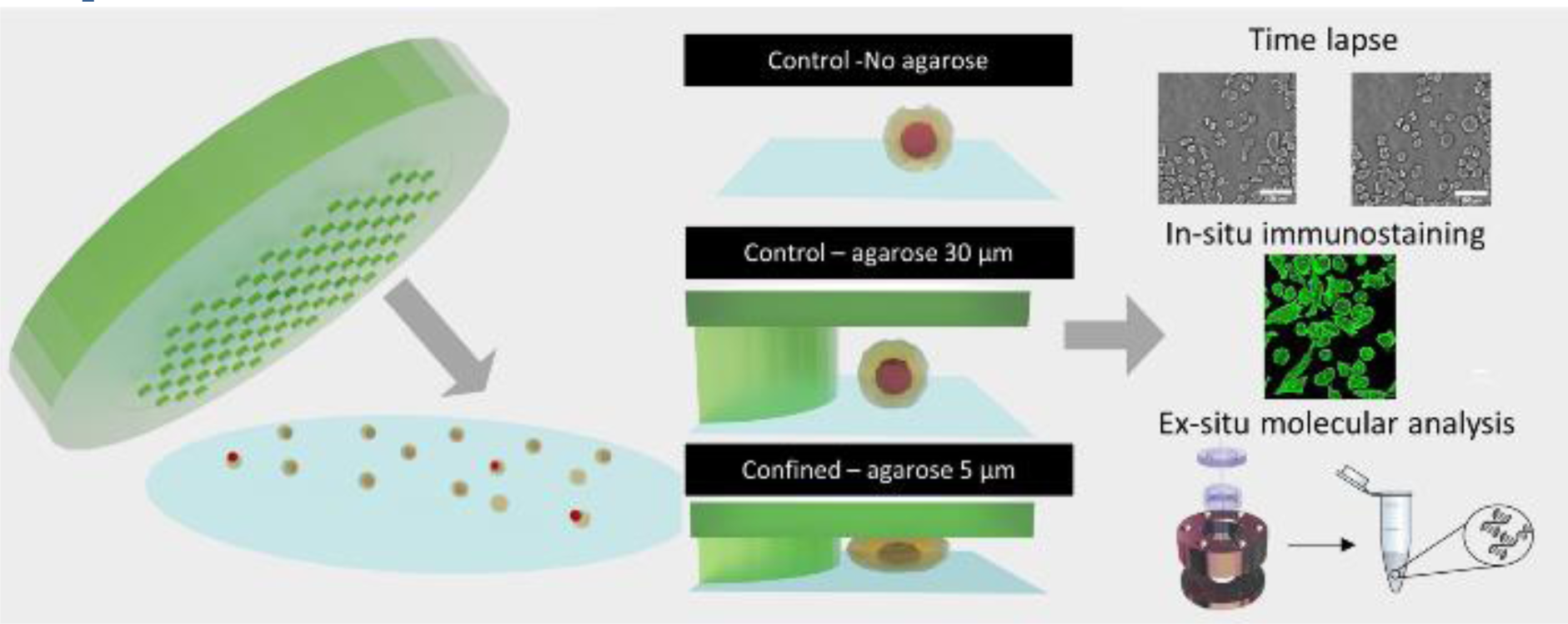

A unique tool to analyze the role of long-term effect of mechanical confinement in normal and pathological conditions

## Introduction

Emerging evidence suggests the importance of mechanical stimuli in normal and pathological situations for the control of many critical cellular functions^1,2^. It has been shown that biomechanical stimuli can induce changes in gene expression^3^, influence stem cell differentiation^4,5^ and are altered in several human diseases such as aging^6^ and cancer^7,8^. In the latter case, the transition of cells towards a cancerous phenotype is accompanied by various mechanical modifications such as ExtraCellular Matrix (ECM) stiffening^9^, increase in interstitial fluid pressure^10,11^, and compressive stress resulting from cell proliferation in a confined environment^12^. In turn, such lateral compression imposed by the surrounding environment strongly drives cancer cells to evolve towards a more invasive phenotype^13^, and is accompanied by changes in gene expression^14^.

Interestingly, while the effect of matrix stiffness is extensively studied in the context of tumor progression^15,16^, stem cell differentiation^17–19^ and aging^6^, few studies have focused on the role of mechanical stresses^20^. This field of research remains underdeveloped due to lack of standard *in vitro* assays enabling quantification of phenotypic and genotypic modifications of cells upon extended mechanical stimulation.

Different polydimethylsiloxane (PDMS)-based microfluidic systems were recently designed to determine the impact of confinement on cell migration^21–24^ and nucleus deformation^25–28^. Using such systems, a switch from a mesenchymal to an amoeboid mode of migration upon cell confinement was highlighted for various mesenchymal cell types^29^ including embryonic progenitor cells^30^. Tunable microsystems enabling the control and analysis of this transition should thus pave the way for understanding the impact of mechanical stress in normal and pathological situations^31^. The ability of cells not only to deform (both their overall and nucleus shape)^32^ but also to repair their nuclear envelope after rupture during migration through confined environments were also evidenced using PDMS-based microsystems^33^.

Nevertheless, drawbacks of such microsystems are two-fold: (1) the rigidity of PDMS (in the MPa range) is several orders of magnitude larger than the rigidity encountered *in vivo* (100-1000 Pa range^7^), and (2) PDMS is impermeable to small water-soluble molecules, leading to fast-medium conditioning if continuous flow is not provided (due to depletion of nutrients or increase in cell secreted-factors). It was our case in early studies on 3-days confinement of various cell lines (**Fig SI.1**). Because these PDMS-based systems could not adequately decouple mechanical signals from other biochemical cues in long-term experiments, they have so far mainly been used to decipher early cell response to mechanical confinement (several hours at most). It is thus crucial to develop new tools for studying long-term confinement, without introducing adverse side effects.

In parallel to PDMS-based microsystems, different hydrogel-based approaches were also developed^34^, including silk^35^, alginate^36^, polyacrylamide^37^ or Poly(ethylene glycol) diacrylate(PEGDA)-derived microsystems^38^. Among them, agarose was uniquely shown to cover a wide range of physiological stiffness, achieved by tuning the type and percentage of agarose^39^. The agarose matrix allows the free diffusion of salt and small molecules (size < 30 nm in 2% agarose^40^, which is the case for most proteins), ensuring passive medium renewal. While it is currently widely used in tissue engineering^39^, it has however only been implemented in microfluidic systems by a limited number of groups^41–43^. The main limitation for its wide-use in lab-on-chip applications is its difficult integration in user-friendly protocols enabling easy sealing and cell recovery. Indeed, as hydrogels are mainly composed of water, various leakage issues remains to be addressed.

Here, we have developed an innovative integrated agarose-based microsystem with a rigidity close to physiological conditions and enabling passive medium renewal. The system mimics the confined state of cells proliferating in a defined volume. The set-up is highly flexible and compatible with time-lapse microscopy, *in situ* immunostaining, as well as standard molecular analyses (qPCR, Western blot). Using this set-up, it was possible to confine, even simultaneously, different cell lines for several days with no major impact on cell viability. Hence, the soft cell confiner described in this manuscript appears as a powerful tool that could be of major interest to address key biological questions in the growing field of mechanobiology.

## Materials and Methods

### Wafer fabrication

A standard photolithography process was used to create the different wafers needed for agarose molding. A thin layer of SU8 photoresist resin (SU8 2000 series, Microchem) was spin-coated onto a silicon wafer and heated on a hot plate. According to the manufacturer’s application notes, the type of resin, the parameters of spin coater and the parameters of baking (**Table SI 1**) were chosen to produce the desired layer height (fixing the height of agarose pillars after agarose molding). The negative photoresist layer was then exposed to UV light through a photomask drawn with Clewin4 (WIEWEB software) presenting the pattern shown in **Fig. 1 A1**: pillars of 440 μm diameter regularly distributed and surrounded by a band of 2 mm to stabilize the structure. After development (with Propylene glycol monomethyl ether acetate −PGMEA 484431, Sigma), the resin that had not been insulated was removed and the wafer was baked on a hot plate to allow proper adhesion of the resin onto the substrate. Finally, the wafer was washed with isopropanol and distilled water. The height of the molded wafer was controlled using a surface profiler (Veeco Dektak 150, contact stylus profilometry techniques).

**Fig 1.**
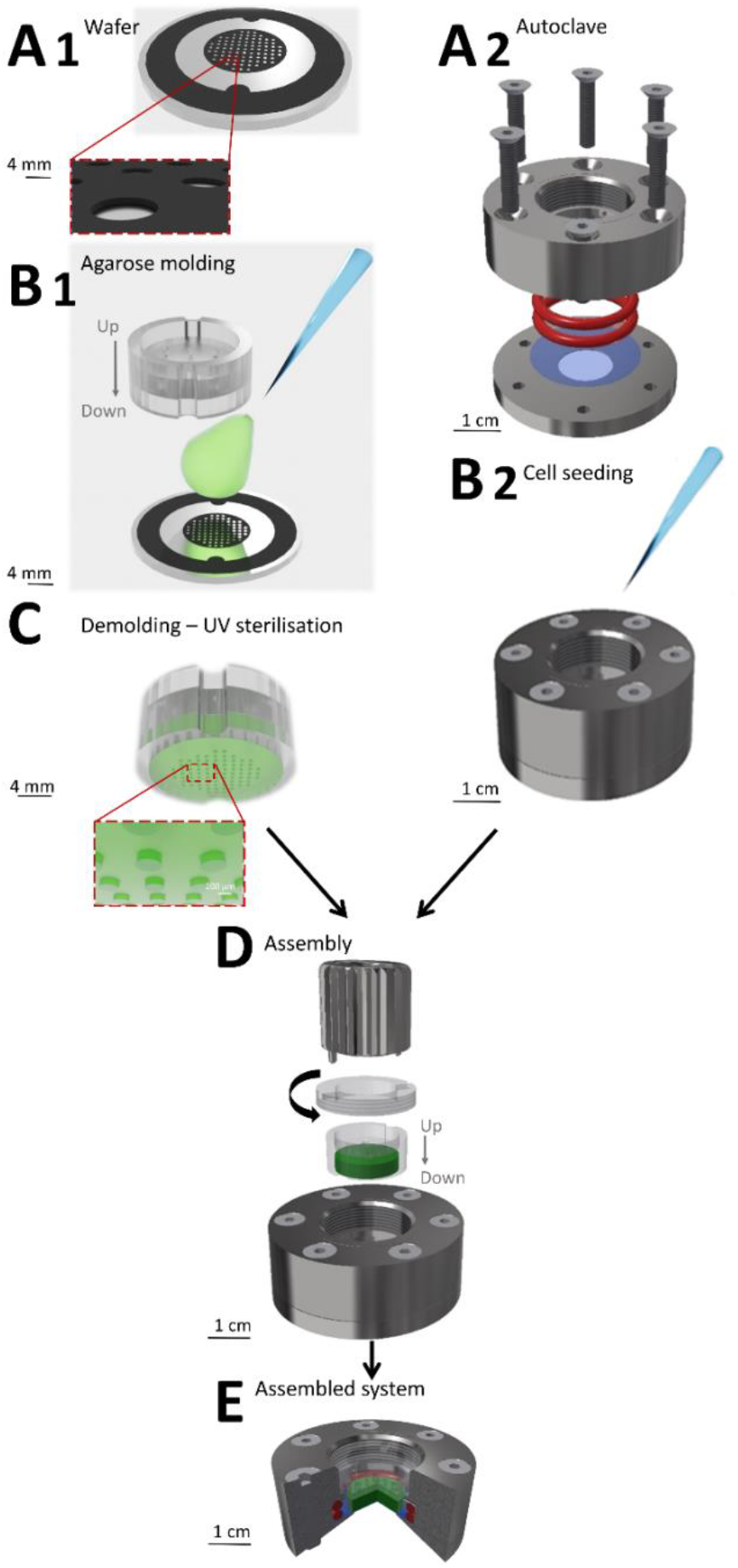
Preparation and assembly of the soft cell confiner. The steps (A1-B1 and A2-B2) of the microsystem preparation can be performed in parallel. **(A1)** Wafer used for agarose molding. **(B1)** Agarose molding on the wafer in the PC holder. **(A2)** Assembly of the system before autoclaving: two pieces of stainless steel hold two o-ring seals (red) and a glass coverslip (blue) **(B2)** Cell seeding onto the glass coverslip in the system **(C)** Schematic representation of the molded gel, which is UV-sterilized before assembly. **(D)** Assembly of the system: PC holder containing the molded agarose gel (green) is placed on top of the seeded cells using a clamping washer tightened with a specific clamping tool **(E)** Cross-section of the assembled system (red: ring seals, blue: glass coverslip, green: agarose gel presenting the pillar network).

### Agarose molding

A solution of standard agarose (3810, Cart ROTH) diluted in distilled water was prepared through a first step of autoclaving at 120°C (4% (w/v)) for 15 min. To visualize the pillars by confocal microscopy, 250 μL fluorescent microspheres (Ø 0.40 μm, BZ5400, Interchim Fluoprobes) sonicated for 30 s were added to 4 mL of agarose solution. The prepared agarose solution (800 μL) was deposited on the pre-warmed wafer molds (placed on a hot plate at 78°C, **Fig. 1 B1**). The polycarbonate (PC) holder was then immediately placed onto the melted agarose and together with the wafer, they were removed from the hot plate and left to set at room temperature (RT) for 10 min. The PC holder was then gently removed from the wafer. Evenly distributed holes were drilled into the gel using a 20 G puncher and through holes present on the PC mold. The holder containing the molded agarose gel presented in **Fig. 1C** was then placed in sterile PBS (Gibco) and sterilized under UV light (20 min each side, Vilber Loumat, 24W, 365/254 nm). The molded agarose was stored in its PC holder at 4°C in PBS until further use. It was replaced by culture medium and incubated at 37°C at least 3 h before mechanical sealing.

### Soft cell confiner assembly

Reversible mechanical sealing between a glass coverslip and the molded agarose was ensured using a custom-made stainless steel system (**Fig. 1 A2**), adapted to the PC holder. Prior to mounting, a glass coverslip (Ø30 mm No1, 631-1585, VWR) and the stainless steel parts were cleaned with 70% ethanol, rinse with distilled water and air dried. The glass coverslip was then placed between the lower and upper stainless steel parts (**Fig. 1 A2**). Sealing was ensured via two silicone o-ring seals (24.50×3.00 mm silicone 70 shores FDA, Fishop) stacked on top of each other in the upper stainless steel part of the system. They were then screwed together before sterilization in an autoclave for 7 min at 134 °C. Screws were tightened gradually and simultaneously in a cross pattern.

### Cell lines and cell culture

The hematopoietic TF1 cell line was obtained and validated as described in^44,45^. Parental TF1-GFP and leukemic BCR-ABL transformed TF1-BA cells were cultured in suspension in RPMI1640 (Gibco) containing 10% fetal calf serum (FCS, Life). ML2 leukemic cells (acute myelomonocytic leukemia) were obtained from F. Mazurier (University of Tours, France) and cultured in RPMI1640 containing 10% FCS. The adherent HS27A cell line, model of mesenchymal stromal cells, was obtained from the ATCC and cultured in RPMI1640 containing 10% FCS. MCF10A cells were purchased from the ATCC and cultured according to recommendations in phenol red-free Dulbecco’s modified Eagle’s medium (DMEM)/F-12 nutrient mix supplemented with 5% horse serum (Life), 10 μg/mL insulin, 0.5 μg/mL hydrocortisone, 100 ng/mL cholera toxin, 20 ng/mL EGF (Sigma), 1% penicillin/streptomycin (Life Technologies).

### Cell seeding and soft confiner mounting

MCF10A and HS27A cells were seeded in the systems overnight before soft confiner mounting (500 μL of a cell solution at 2×10^5^ cells/mL). For TF1 cells, a fibronectin solution (F.895 Sigma-Aldrich, 50 μg/mL in NaHC0_3_) was first used to coat the glass surface (30 min incubation in the system at 37°C). Excess fibronectin was removed through three washes. Cells were then seeded onto the glass coverslip (5.6×10^5^ cells/mL for TF1-GFP, 10.2×10^5^ cells/mL for TF1-BA, 500 μL /system), and incubated for 2 h at 37°C to allow proper adhesion to the substrate. Concerning the co-culture experiment, HS27A cells were seeded at 2.5×10^5^ cells/coverslip and then incubated for 24 h. ML2 cells were then added (3×10^5^ cells/coverslip) and incubated for 2 h at 37°C.

After cell adhesion, the seeded cells were gently washed three times to replace the medium with pre-warmed fresh culture medium (500 μL, **Fig. 1 B2**). The PC mold containing agarose gel was placed in the system and a clamping washer was tightened with a specific clamping tool (**Fig. 1 D** and **SI movie 1**). The gel was then tightly in contact with the glass coverslip supporting the cells. A reservoir of 500 μL of culture medium was added above the PC holder and incubated for 1 h at 37°C. The molded agarose was then washed three times (5 min each) with pre-warmed culture medium.

To assess that neither the stainless steel assembly nor the PC-holder with molded agarose affected cell behavior, 2 control conditions were used for each experiment:

1. cells on a glass coverslip in the stainless steel assembly, with no molded agarose (1000 μL culture medium). This will hereafter be referred to as Control throughout the manuscript.
2. cells on a glass coverslip in the stainless steel assembly with agarose molded with an array of pillars of 30 μm in height, larger than the height of the cell population investigated (500 μL of culture medium in the molded agarose + 500 μL above the PC holder). This will hereafter be referred to as 30-μm throughout the manuscript.

### Immunostaining within soft cell confiner

After confinement, cells were fixed *in situ* with 4% paraformaldehyde (PFA, 15714, EM Grade): the cell culture medium was removed and the samples were washed three times with PBS. 4% PFA was then added and incubated for 20 min at RT. After incubation, the samples were washed three times with PBS and incubated with 0.5% Triton X-100 (2156825000, Acros organics) in PBS for 10 min at RT for permeabilization followed by three consecutive washes with 0.1% Triton X-100 every 5 min. After permeabilization, samples were blocked with 3% BSA (A2163, Sigma) 0.1% Triton X-100 in PBS for 20 min at RT to inhibit non-specific binding of antibodies. Cells were initially incubated with Alexa 546 Phalloidin (A22282, Thermofisher, 1:50 in 0.1 % T-X 100 in PBS) for 20 min at RT and washed three times with PBS. Samples were finally incubated with NucGreen^™^ (Thermofisher, R37109, 2 drops/mL in PBS) for 15 min and washed three times with PBS.

### Cell viability

Fresh culture medium was added in each system every 2 days. To assess cell viability, agarose gel was dismounted and cell viability was monitored with calcein (Thermofisher) and propidium iodide (PI, Sigma) labelling. Calcein labels the cytoplasm of viable cells in green, PI labels nuclei of dead cells in red. A maximum volume of culture medium was removed and cells were then washed once with pre-warmed PBS. Calcein (1 μM) and PI (2 mg/mL) diluted in pre-warmed sterile PBS were then incubated for 20 min at 37°C before epifluorescence microscope analysis.

Control live and dead cells were tested in parallel. For live cells, cells in classical 2D cultures were incubated for 24 h at 37°C. For dead cells, 70% ethanol was added 30 min prior to staining (**Fig. SI 2**).

### Microscopy

Cells were observed with inverted microscopes (Leica DM IRB or Leica DMI8) using phase-contrast imaging. A 10x objective was used to follow cell proliferation and a 5x objective to observe their viability with calcein/PI staining. Images were taken every 24 h. In parallel, phase-contrast time-lapse imaging was performed for 24 h in a controlled (C0_2_, temperature and humidity) environment. A motorized x-y stage enabled the concomitant recording of up to 10 regions for each system every one or two hours (**SI movie 2 for TF1-BA and SI movie 3 for MCF10A**).

Fixed and co-cultured cells were visualized using a Leica SP5 confocal microscope or a Zeiss LSM 880 confocal microscope with a 20x dry objective (NA 0.65). Z-stacks of live cells in the soft confiner were also acquired at 20x magnification (dz= 0.4 μm for each stack).

### Image analysis and quantification

In order to monitor the proliferation of different cell types, the number of cells for each image was determined with the free program ImageJ using a custom-written routine based on the Find maxima tool of the software. The cell density (in cells/cm^2^) was analyzed for 10 different positions for each tested condition. At least three samples and two independent series of experiments were performed for each cell type. The experiment was stopped when confluence was reached (day 3 for HS27-A, day 2 for TF1-BA, day 1 for MCF10A).

Area of TF1-GFP live cells, area and circularity of TF1-BA fixed cells were assessed with a homemade Matlab program. Images were filtered using a Wiener adaptive filter and cells were then separated from the background using threshold detection and converted to binary images. Sequential steps of morphological reconstruction were performed and cells were individually detected based on different parameters (distance between centroid, area, eccentricity). Four different areas within the soft confiner were analyzed for each condition.

Nucleus circularity was defined by:

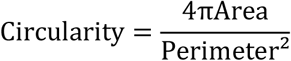

### Western blot

Western blot analysis was performed on HS27A cells confined during 3 days, proteins were then extracted using RIPA buffer. Per lane, 15 μg of proteins were loaded onto gels prior to conducting SDS-PAGE and transferring onto polyvinylidene difluoride membranes (Bio-Rad). Membranes were then incubated with monoclonal antibodies against Cyclin B1 (PC133, Calbiochem) and GAPDH (#8884, Cell Signalling Technology). Specific binding of antibodies was detected using appropriate secondary antibodies conjugated to horseradish peroxidase, and visualized with Clarity Western ECL Substrate (Bio-rad), on ChemiDoc Gel Imaging system (Bio-rad). Densitometric analyses of immunoblots were performed using ImageJ.

### Flow cytometry

Flow cytometry cell sorting experiments were carried out using HS27A-Turquoise and ML2-Cherry. Briefly, the lentivirus expressing mTurquoise2-Tubulin was constructed by cloning the mTurquoise2-Tubulin sequence from pmTurquoise2-Tubulin (a gift from Gadella Dorus, Addgene plasmid #36202^46^) into the CSII-EF-MCS vector. Lentiviruses were produced in 293T cells by transfecting lentiviruses with the helper plasmids pMD2.G and psPAX2 (a gift from D. Trono, Addgene plasmids #12259 and #12260), following Addgene’s instructions. Recipient cells were infected at a low multiplicity of infection (moi < 1) and finally sorted on a BD FACSAria III SORP cell sorter.

After 3 days confinement, the HS27A-Turquoise and ML2-Cherry cells were analyzed using the BD LSRFortessa cell analyzer.

### Statistical analysis

Data were expressed as mean ± standard deviation. The statistical significance of differences between conditions was analyzed using the Prism software (GraphPad). The Mann-Whitney test was used for figs. 3A, 3C, 4A, 6A and unpaired t-test for figs. 2D, 4D, 4E, 5D. Differences with a p-value under 0.05 were considered statistically significant. The significance is indicated by asterisks in figures (* p < 0.05; ** p < 0.01, *** p < 0.005).

**Fig 2.**
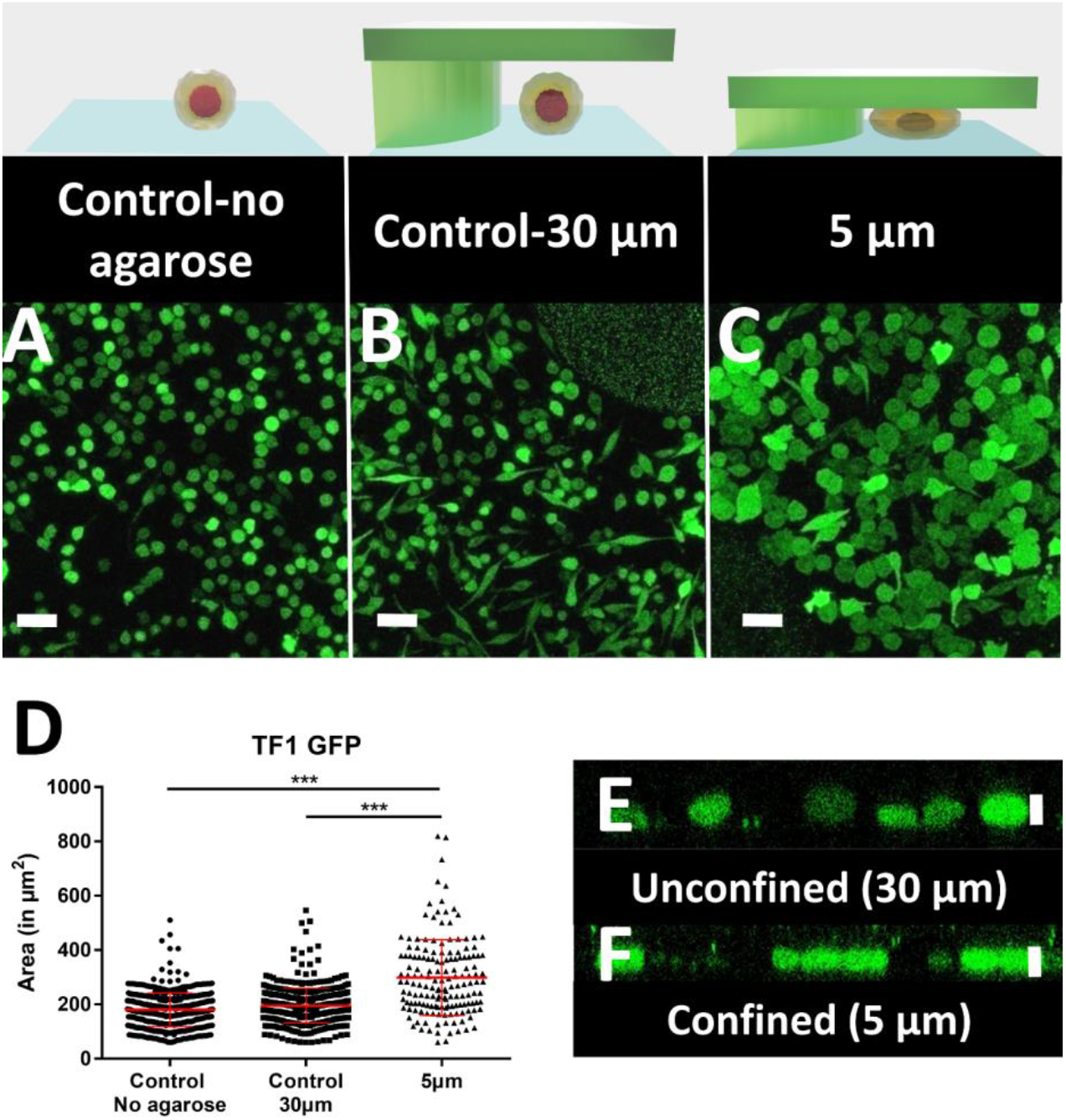
Quantification of cell morphology under confinement. **(A-C):** Morphology of immature TF1-GFP hematopoietic cells for control **(A)** and for 30-μm and 5-μm (B and C, respectively). Scale bar = 20 μm. **(D)** Quantification of projected area by automatic image analysis. Projected area was similar for control and 30-μm, while it significantly increased for 5-μm confinement (Unpaired t-test n = 160 at least for each condition) **(E-F):** z-section of unconfined **(E)** and confined **(F)** cells. Vertical scale bar = 5 μm.

## Results

As our previous attempts at setting-up a reliable cell confinement tool using a PDMS-based confiner similar to^28^ fell short of our expectations, we were committed to developing a hydrogel-based microsystem in which culture conditions could be fully controlled (**Fig. 1**). However, the implementation of a reliable and reproducible cell confiner using a hydrogel-based microsystem is not trivial. The hydrogel has to be properly sealed on the glass coverslip, without compressing the gel or inducing leakage. We achieved these objectives in an ultimate user-friendly design (**Fig. 1**). The soft cell confiner was designed to enable the concomitant production of agarose molds (**Fig. 1 A1-B1**) and the mounting of coverslips in an autoclavable stainless steel chamber (**Fig. 1 A2-B2**). A plastic holder was specially designed to improve agarose molding and handling (**Fig.1 C**). After cell seeding onto the coverslip, the plastic holder and a dedicated clamping washer and clamping tool enabled the reproducible and rapid mechanical sealing of the molded agarose onto the seeded cells (**Fig.1 D, E and movie 1**). Owing to this system, removal of the molded agarose and retrieval the confined cell population at the end of a long-term experiment was relatively rapid and simple.

Evenly distributed holes in the plastic holder enabled the renewal of culture medium through the upper part of the plastic holder without disturbing confinement conditions or cells. The efficiency of medium renewal was checked by analyzing the evolution of fluorescent intensities upon medium exchange (**Fig. SI 3**). A fluorescent medium placed in the upper part reaches the lower part (where the seeded cells are) by pure diffusion over a characteristic time-course of 7h30.

We first assessed that cells were properly and homogeneously confined using the non-adherent hematopoietic cell line TF1-GFP (**Fig. 2**). While cells cultured for 1 day displayed similar size and morphology for control conditions and 30 μm pillars (**Fig. 2A-B**), the area of cells under 5-μm confinement was much larger (**Fig. 2C**). The projected area increased from 196 ± 3 μm^2^ to 300 ± 11 μm^2^ under 5μm-confinement (**Fig. 2D**), while the height was restrained by the pillars (**Fig. 2E vs 2F**).

In order to fully validate this confining device, we investigated the confinement of the stromal cell line HS27A. After 3 days, while HS27A proliferation inside the PDMS-based microsystem was impacted even with the 30 μm pillars (**Fig. SI 1**), in the same condition the hydrogel-based microsystem had no impact on cell proliferation (**Fig. 3A** control *vs* 30-μm, no significant difference). The significant decrease in cell proliferation induced upon 5-μm confinement (**Fig. 3A**) can hence be interpreted as the mechanical cell response to the imposed 5-μm confinement applied for 3 days. The confined cells were then harvested after 3 days and further processed for protein analysis by Western blot. The level of cyclin B1 was similar for control and 30 μm pillars (**Fig. 3B**), confirming the lack of impact of the confining chamber components on proliferation. Conversely, we observed a strong decrease in cyclin B1 in the 5-μm confinement condition (**Fig. 3B** 30-μm *vs* 5 μm), confirming a decrease in cell proliferation for this stromal cell line under confinement.

**Fig 3:**
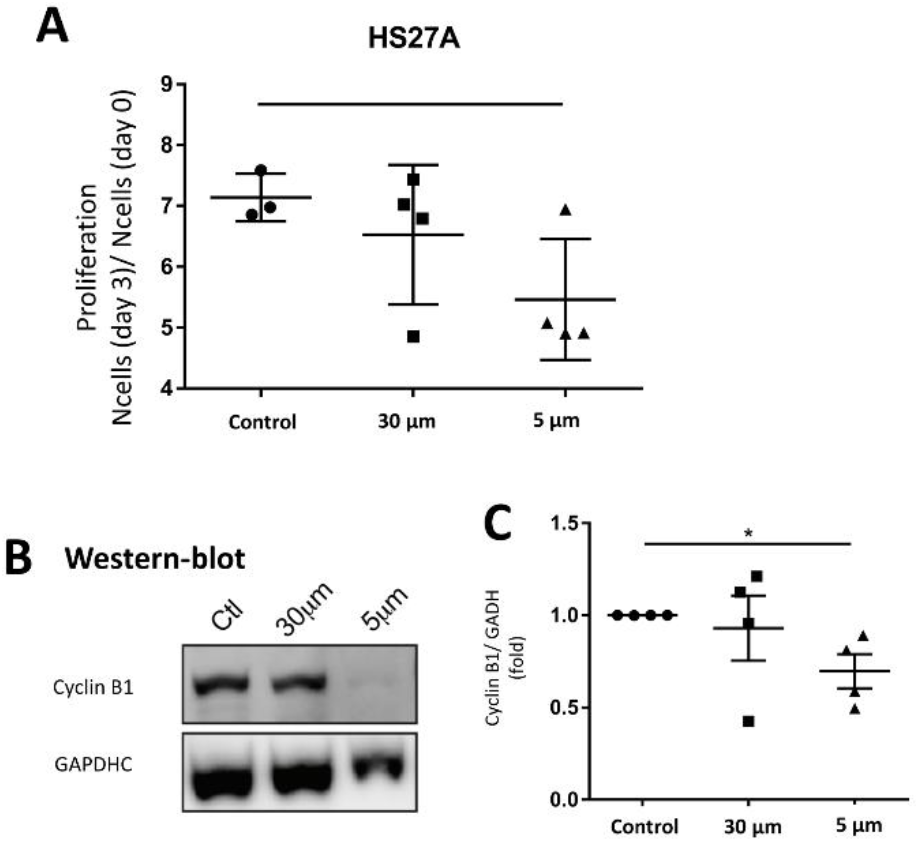
Proliferation of HS27A cells. **(A)** Bar graph showing the proliferation ratio of stromal cells HS27A over 3 days for control and for 30-μm and 5-μm. Density was analyzed for 10 positions for each sample, n = 4 (Mann-Whitney test n.s. not significant) **(B)** Western blots showing CyclinB1 levels from HS27A cells for the three conditions **(C)** Bar graph showing CyclinB1 level (GAPDH used as internal control) n = 4 for the three conditions.

To demonstrate the versatility of this soft cell confiner, it was then challenged by culturing the hematopoietic leukemic cell line, TF1-BA. Indeed, as these cells are poorly adhesive, even in the presence of fibronectin coating, analysis on this confinement system is experimentally challenging. No significant difference in cell proliferation was induced by the microsystem even after 2 days of confinement (**Fig. 4A** control *vs* 30-μm condition), reinforcing previous results obtained with the HS27A cell line. In addition, despite their transformation, these leukemic cells appeared to be sensitive to mechanical stress as we measured a significant decrease in cell proliferation upon 2 days of confinement when comparing 30-μm and 5-μm conditions (**Fig. 4A**, 1.9 +/- 0.2 fold increase in cell number for 5-μm compared to 3.40 +/- 0.4 fold increase for 30-μm).

**Fig. 4.**
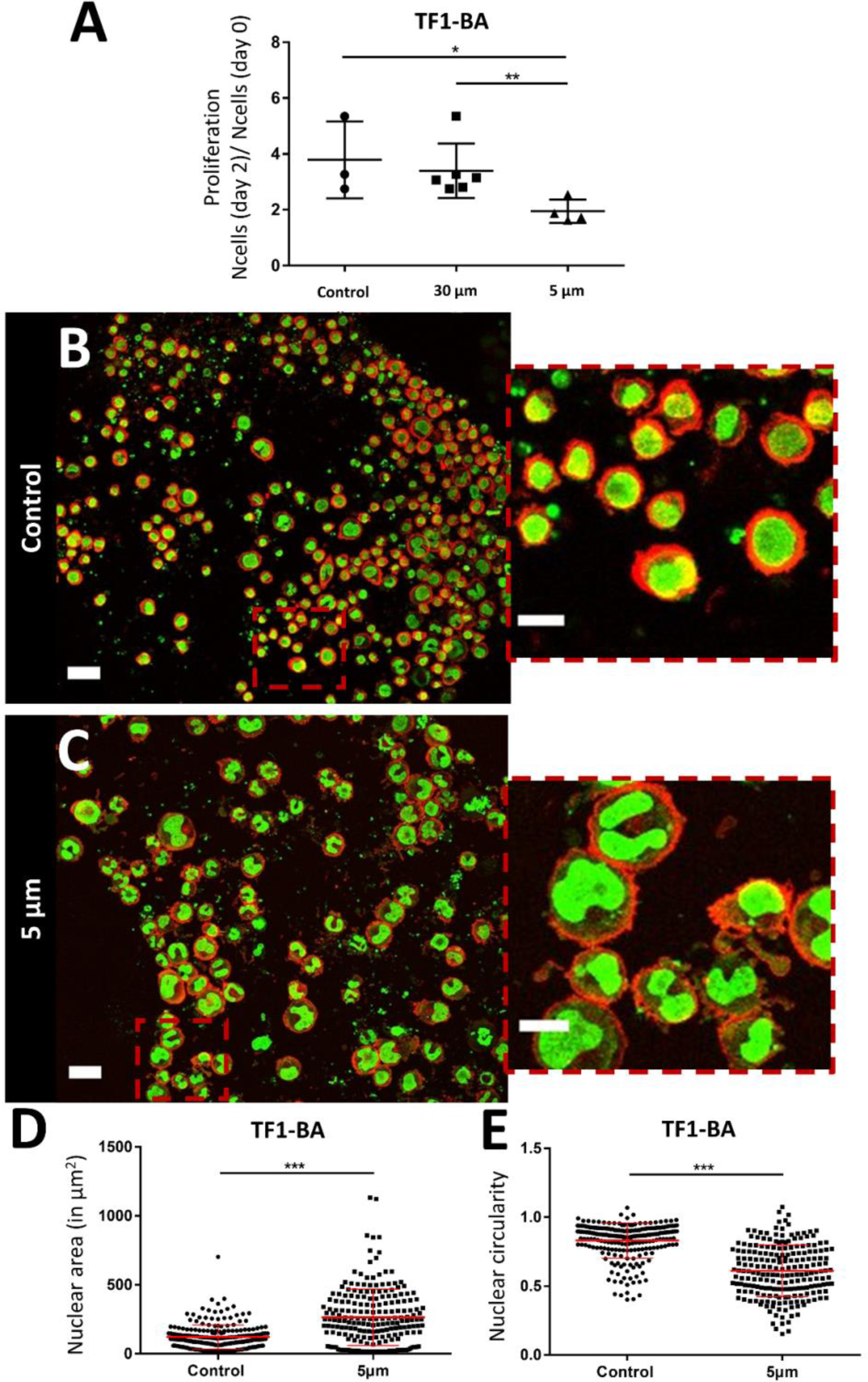
Proliferation of TF1-BA and in situ immunostaining. **(A)** Bar graph showing the ratio of proliferation of TF1-BA immature hematopoietic cells over 2 days for control, and under molded agarose with pillars of 30- and 5-μm. We observe a significant decrease in proliferation under confinement compared to controls (at least 2 independent experiments, performed at least in triplicate for each condition and density were analyzed on 10 positions for each sample). **(B-C)** In situ Immunostaining of TF1-BA cells after 1 day in the soft cell confiner for control **(B)** or under 5-μm confinement **(C).** Left scale bar = 50 μm, right scale bar = 20 μm. Actin (phalloidin) in red and nuclei in green. **(D-E)** Quantification of nuclear circularity and nuclear area of TF1-BA cells for control with no agarose and under confinement (5-μm) after 1 day in the soft cell confiner (confined cells present a larger area and deformation of their nucleus compared to control). At least 213 nuclei were analyzed per condition.

Our soft cell confiner is not only compatible with live imaging but also enables *in situ* immunostaining under confinement. We stained both nuclei and actin under confinement (**Fig 4B-C**). The nuclear projected area appeared to be larger upon confinement (**Fig 4D**), as indicated by an increase in the mean projected nucleus area from 122 ± 6 μm^2^ for control, up to 265 ± 14 μm^2^ for 5-μm confined cells. In addition, we observed that many nuclei were highly deformed, exhibiting a non-circular or polylobed shape (**Fig 4E**, decrease of mean nuclei circularity from 0.83 ± 0.01 to 0.61 ± 0.01).

Our soft cell confiner is also compatible with more complex cell population analysis, mimicking multiparametric and heterogeneous cellular microenvironments, such as stromal cell interactions. To illustrate such complex cell interactions, adherent stromal HS27A cells were co-cultured with the suspended hematopoietic ML2 cells and analyzed in the soft cell confiner system using a confined height of 9-μm to take into account the two layers of cells (**Fig. 5A**). In this setting, we used multicolor labeling of the different cell lines to distinguish them *in situ* and to measure their area upon confinement (**Fig 5A**). This confirmed that despite the increased complexity and heterogeneity in cell population, the system still enabled the measurement of cell area by confocal microscopy. Here, we observed an increase in the mean projected hematopoietic cell area upon confinement (**Fig. 5B-C**) from 216 ± 133 μm^2^ for 30-μm to 329 ± 155 μm^2^ for 9-μm (**Fig. 5 D**). In addition, our system allowed us to recover viable cells after 3 days of confinement to perform various functional assays, such as cell sorting by flow cytometry (**Fig. 5 E**).

**Fig. 5:**
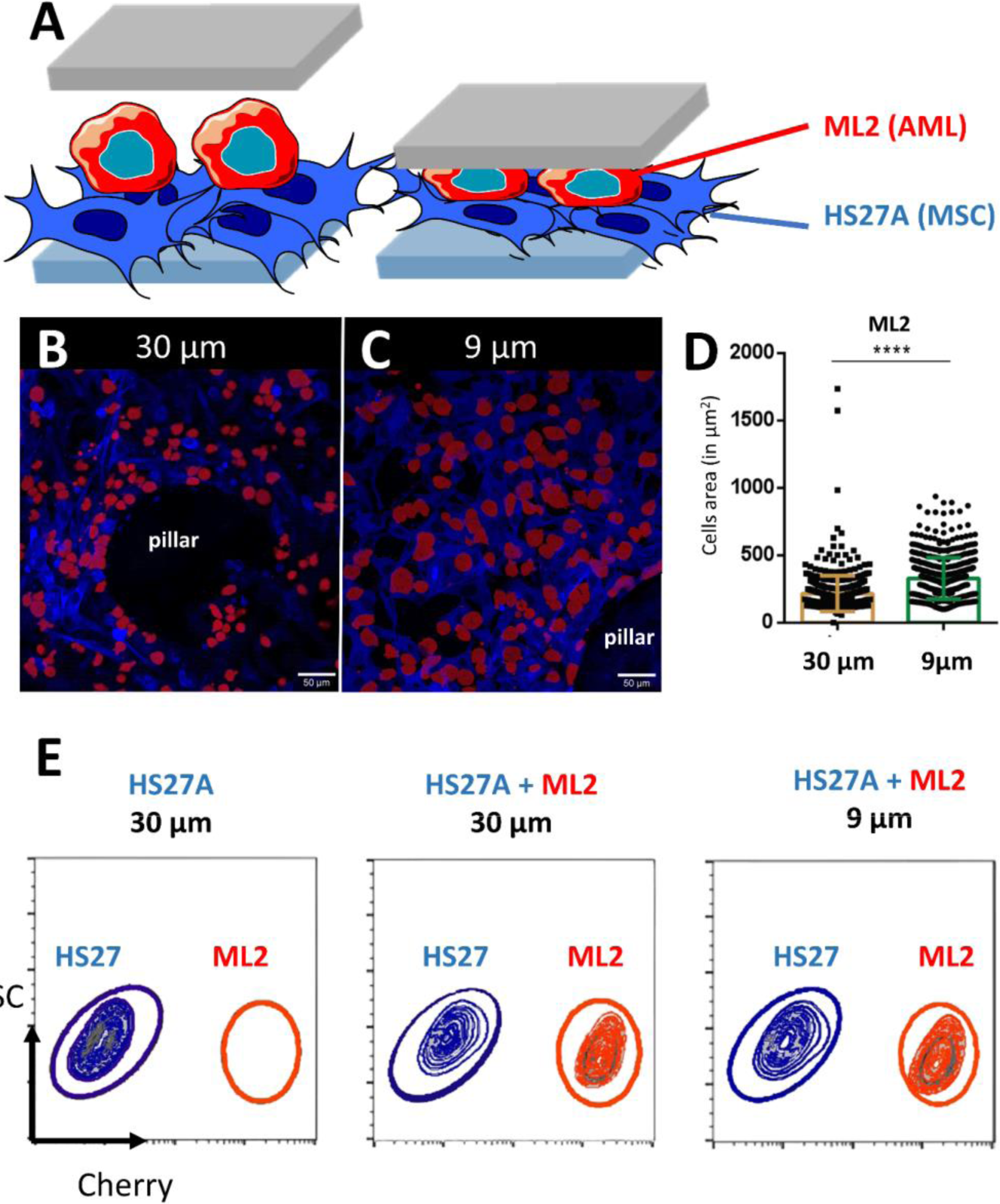
Set-up mimicking the complexity of the tumor microenvironment. **(A)** Schematic representation of the two layers of cells inside the soft cell confiner. **(B-C)** Representative confocal images of leukemic cells (ML2, red) seeded on top of stroma cells (HS27A, blue) and inserted into the confiner with a height greater than both layers (h = 30-μm, unconfined, B) or smaller (h = 9-μm, confined, C) for 3 days. Scale bar = 50 μm **(D)** Bar graph showing ML2 cell area quantification from unconfined (A, the average projected area of 216 ± 133 μm^2^) or confined (B, the average projected area of 329 ± 155 μm^2^). **(E)** Representative FACS plots following 3 days confinement in co-culture analyzed for cell content in HS27A-Turquoise and ML2-Cherry.

Finally, the compatibility of the soft cell confiner with long-term experiments was assessed using an epithelial breast cell line, the adherent MCF10A cells. This immature mammary stem cell line displays contact inhibition of proliferation and can thus be cultured in high-density conditions for several days. Once again, we verified that the hydrogel-based microsystem itself had no impact on cell proliferation (**Fig 6A**, no significant difference in proliferation ratio for control and 30-μm after 1 day-proliferation ratio of 1.8+/- 0.1 and 1.9 +/- 0.1, respectively-). Consistently with our previous findings, a significant decrease in cell proliferation was observed under 5-μm confinement (**Fig 6A**), with the proliferation ratio dropping below a significantly lower value than the two controls (proliferation ratio of 1.5 +/- 0.1 after 1 day). Finally, we confirmed that most cells, except for the ones under pillars, were alive (**Fig SI 4, live/dead staining after 1 day**). Of note, it was possible to culture MCF10A cells under confined conditions for up to 8 days without induction of any significant effect on cell viability, as assessed by live and dead cell staining (**Fig. 6B-C and Fig SI 2** for control of staining).

**Fig 6.**
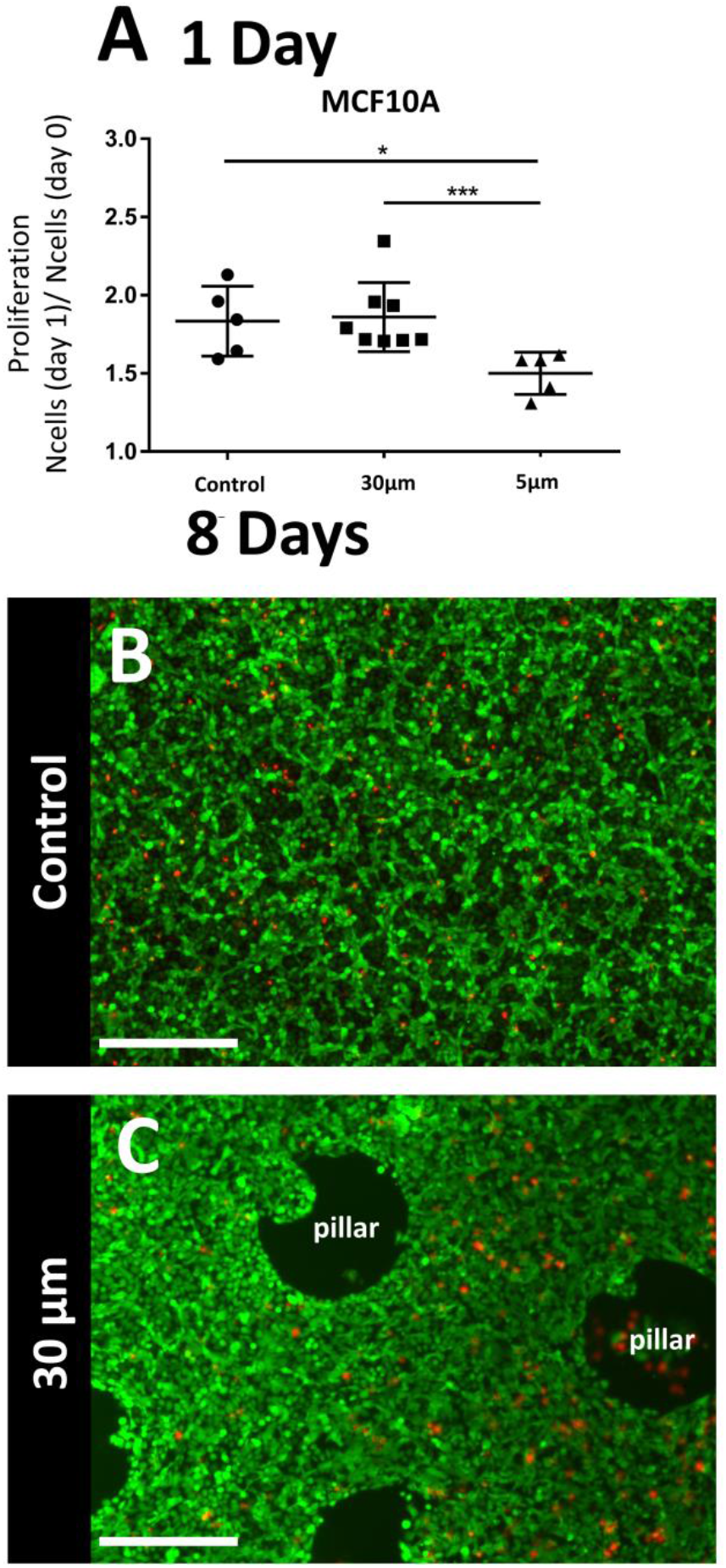
Long-term analysis of epithelial cell lines with contact inhibition under confinement. **(A)** Bar graph showing the proliferation ratio of epithelial MCF10A cells after 1 day in the soft cell confiner with various confinement conditions. At 5-μm confinement, a significant decrease in proliferation under confinement was observed found (at least 2 independent experiments in duplicate or triplicate for each condition. For each system, the number of cells was assessed in 10 different positions) **(B-C)** Live and dead staining of an 8-day MCF10A confluent monolayer in the soft confiner: control with no confinement **(B)** 30 μm pillars. Green: calcein, red: propidium iodide. Scale bar = 400 μm.

## Discussion

The agarose-based microsystem presented herein defies previous cell confining microsystems. Indeed, using PDMS-based microsystem, it has so far only been possible to monitor early cell response to confinement (several hours at most), due likely to limited access to nutrients. In our soft cell confiner, the porous nature of the confining wall enables easy medium renewal. Cells in a confined state can be cultured for several days without nutrient availability impairing their survival.

Several other alternative set-ups have been developed to apply a defined stress on an entire cell population for prolonged periods of time, by embedding them in agarose^12^ or in extracellular matrix^47^ or using a transmembrane-based pressure device^48^. The drawback of such devices is that they are not compatible with high-resolution microscopy due to the thickness of the gels or the transwell geometry. Only analyses at end-point are possible, limiting all real-time analyses on cellular adaptation to mechanical stress. Here, we have developed a system that combines the advantage of PDMS-based microsystems with a transparent chip geometry on glass coverslips, enabling a real-time dynamic analysis, as well as a long-term monitoring of cells. Our soft cell confiner device integrates different biophysical and biological approaches that were barely achieved with current existing devices. At the end of the experiments, the porous nature of the confining wall and the compatibility of our system with high resolution microscopy provided highly interesting analyses such as *in situ* immunostaining and *in situ* confocal microscopy. Alternatively, the entire cell population could easily be collected and processed using standard molecular biology protocols (qPCR, Western blot) or functional assays.

Because our device does not impact proliferation in the absence of confinement, we can truly decipher the impact of confinement on cell proliferation. In the current study, a decrease in proliferation under spatial confinement was observed for all cell lines tested over one to three days in the confiner. Similar results reported a higher frequency of quiescent cells upon confinement within a stiff 3D matrix^49^. These results also corroborate those obtained using PDMS set-ups^50,51^ and atomic force microscopy (AFM)^52^, where it was reported that confinement delays mitotic progression.

We further analyzed nucleus shape and actin at a defined time-point via immunostaining. Interestingly, for immature TF1-BA hematopoietic cells, after 1 day under 5-μm confinement, nuclear shape and projected area were highly modified. The interplay between actomyosin contractility and nuclear deformation in the regulation of nuclear transport of signaling molecules has been highlighted in the past few years^53,54^. Dynamic studies are now needed to unravel how nuclear deformation and mechanosensitivity modulate gene expression and cell fate spatiotemporally. As our soft cell confiner is compatible with time-lapse microscopy, it could also be used to follow nucleus deformation dynamically for several days, using dedicated fluorescent constructs such as Histone or lamins A/C. We plan to use our soft cell confiner system to investigate these important issues in the near future.

Importantly, we demonstrated that our confiner can be used for both adherent and non-adherent cells. It can also be used to disentangle the interaction between different cell types, as shown in this manuscript with two layers of cells (stromal and leukemic cells). Hence, it could be a valuable tool to analyze the dynamic interplay between heterogeneous cell populations in response to mechanical stress.

Cell response to the application of a local stress has also been extensively investigated using AFM set-ups^52,55,56^. While this single-cell approach technique can provide interesting insight into cell response to the application of a local force, it is relatively low-throughput and requires additional expertise to be properly interpreted. Our soft cell confiner is a complementary approach, as we do not apply a defined force but we impose a defined deformation (set by the height of the pillars).

The confining matrix rigidity can be adjusted to closely match physiological conditions (rigidity of [100 Pa-5 kPa^9,57,58^) by tuning the concentration and the type of agarose (standard, low or ultra-low melting agarose). In addition, to ensure a similar stiffness on both top and bottom walls, the coverslip can be coated with an additional soft agarose layer. The soft cell confiner system can hence be used to decipher the influence not only of confinement but also of matrix stiffness (in combination or separately). Such flexibility could be of primary importance to unravel the role of these two important biochemical cues in various biological contexts.

Of note, fluorescent beads incorporated into the agarose could be used to measure cell-generated forces in response to the imposed confinement by 3D traction force microscopy (TFM,^25,59–62^). If the gel is soft enough to be deformed by the cells, a dynamic quantification of the compression force sensed by the cell could be retrieved. Such quantification is out of the scope of this paper but work is ongoing within our team to determine whether the cells reorganize in response to confinement to decrease the imposed mechanical stress, and if so, at what time scale (hours/days).

In this new set-up, we have made the arbitrary choice to confine cells in the z-direction only. However, it is also possible to confine cells in x,y and z directions using a dedicated design. Indeed, compared to other hydrogels, agarose gels do not swell, and it is thus easily molded in various shapes and aspect ratio.

In the current set-up, the pure agarose confining roof provides no adhesive groups for cell attachment. Nevertheless, it is possible to analyze the effects of adhesion by interpenetrating the network of agarose with collagen^63^, PEGDA with covalently immobilized RGD peptides^64^ or silk^65^. This could offer the possibility to analyze the role of various extracellular matrix proteins on cell response to mechanical confinement.

## Conclusion

The hydrogel-based system detailed in this manuscript paves the way for new approaches and concepts in cell biology.

We anticipate that this device could be a valuable tool for the fundamental understanding of the effect of cell confinement on various hallmarks of cancer progression and resistance, and in particular to decipher the role of nuclear mechanosensitivity. Links between cell resistance to treatment and mechanical stimuli have been highlighted recently^49,66,67^. Such mechanical cues could hence lead to new therapeutic approaches^68^ that could be tested using such systems.

## Contribution of each author

AP: Methodology, Investigation, Validation, Formal Analysis, Visualization, Writing – Original Draft
SL: Investigation, Validation, Visualization, Writing – Review & Editing
HDA: Conceptualization, Writing – Review & Editing
BL: Investigation
GS: Methodology, Visualization
SS: Investigation
FA: Supervision
BG: Resources, Supervision
JPR: Funding Acquisition
SG: Conceptualization, investigation, Supervision, Writing – Review & Editing,
VMS: Conceptualization, Supervision, Writing – Review & Editing, Funding Acquisition, Project Administration
CR: Conceptualization, Supervision, Writing – Original Draft, Writing – Review & Editing Funding Acquisition, Project Administration

## Acknowledgments

This work was supported by the Institut Convergence PLASCAN, the Ligue Contre le cancer foundation, the “Institut Universitaire de France” (IUF), the “Fondation pour la Recherche Medicale” (FRM) for part of A. Prunet’s salary and the “Fondation de France” and Alte SMP for part of S. Lefort’s salary.

We thank Y. Chaix, R. Zagala, M. Jacquemin, Q. Cassar, A. Damn for their help in the early development of this project, as well as L. Barral for LAM/stroma pictures and Q. Ducerf for his help with MCF10A cells.

## Supplementary Data

**Figure SI 1:**
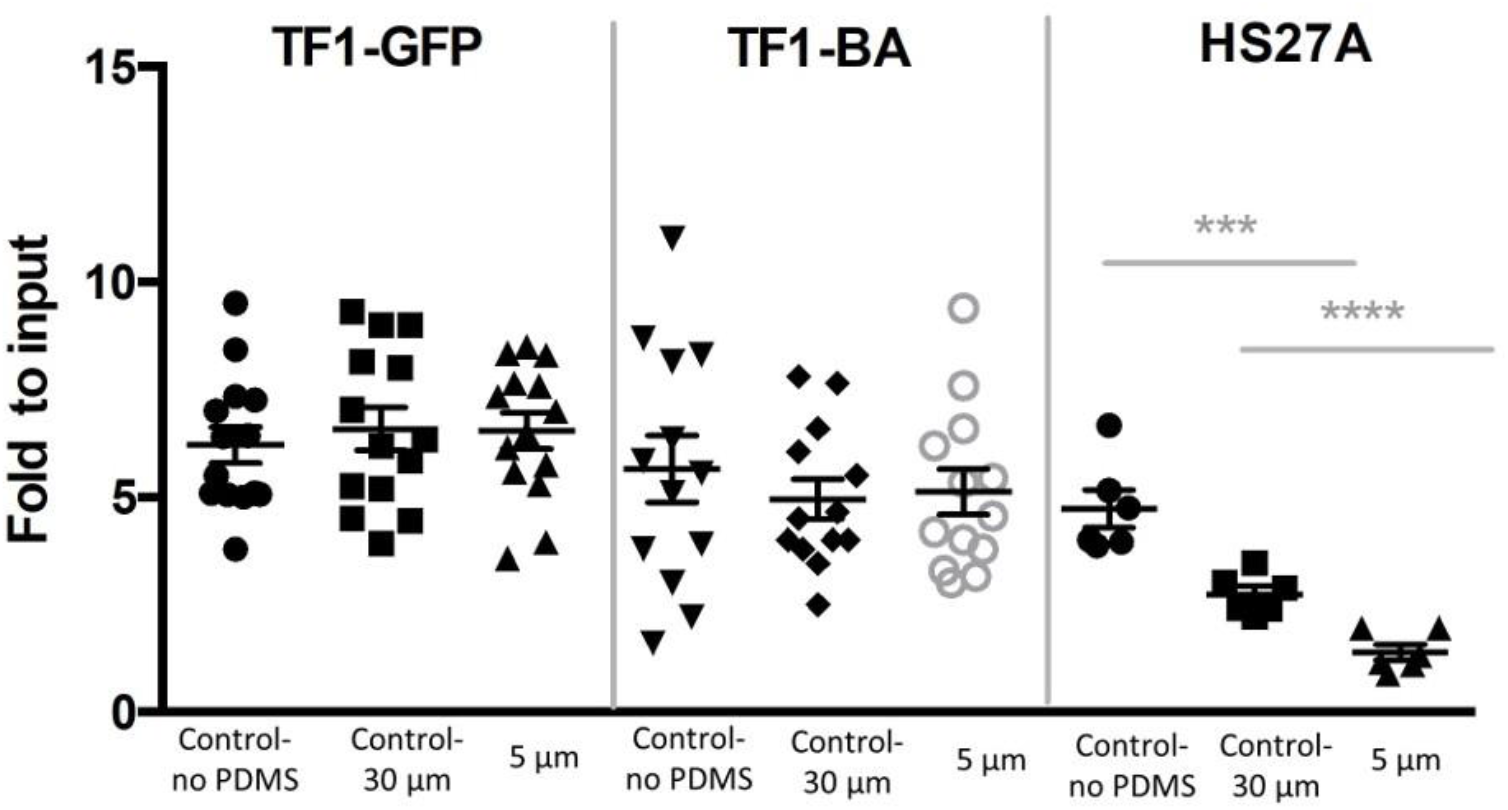
Proliferation of TF1-GFP, TF1-BA and HS27A cells over 3 days in a PDMS-based confiner. In preliminary experiments, the effects of 3 days confinement on hematopoietic cell lines and their stromal cell supports were investigated using a PDMS-based confiner similar to^28^, leading to distinct behavioral changes between cells. Proliferation of the HS27A cell line was impacted by the system itself, since both the 5-μm confinement and the 30-μm control significantly decreased cell proliferation. We thus hypothesized that this effect was likely reflecting the lack of control of medium renewal, and/or hypoxic conditions induced by the presence of the glass coverslip supporting the PDMS pillars. These preliminary results were difficult to interpret as we could not discriminate if the cell response was due to mechanical stress and/or nutrient or oxygen deprivation.

**Table 1.** Parameters used for wafer substrate fabrication: photolithography process. For 9 μm, the process for 5 μm was repeated twice.

| Thickness | Resin used | Spin coater parameter | Soft Bake | Insolation | Post Exposure Bake |
| --- | --- | --- | --- | --- | --- |
| 5 $\mu\text{m}$ | SU8 2005 | Speed 500 rpm,<br>acceleration 200, 10s<br>Speed 2500 rpm,<br>acceleration 500, 30s | 2 min,<br>95°C | 110 mJ | 3 min, 95°C |
| 30 $\mu\text{m}$ | SU8 3050 | Speed 500 rpm,<br>acceleration 1, 10s<br>Speed 4000 rpm,<br>acceleration 3, 30s | 10 min,<br>95°C | 170 mJ | 1 min, 65°C and 5<br>min, 95°C |

**Figure SI 2:**
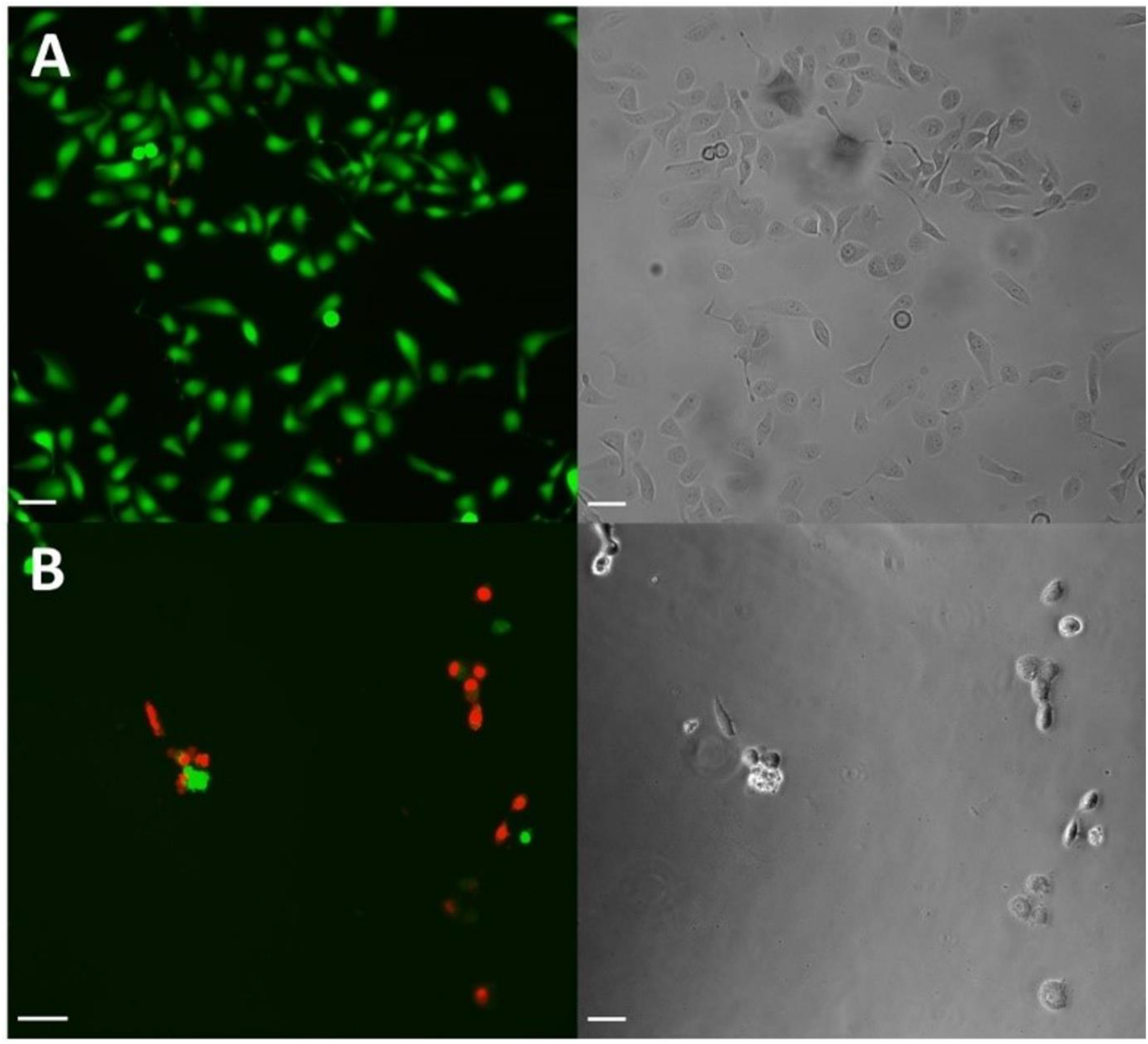
Control of live and dead MCF10A cells on glass coverslips after 1 day of incubation. (A) Control live and (B) control dead cells obtained by adding 70% ethanol for 30 min prior to imaging. Calcein AM is represented in green and propidium iodide in red, scale bar = 100 μm.

**Figure SI 3:**
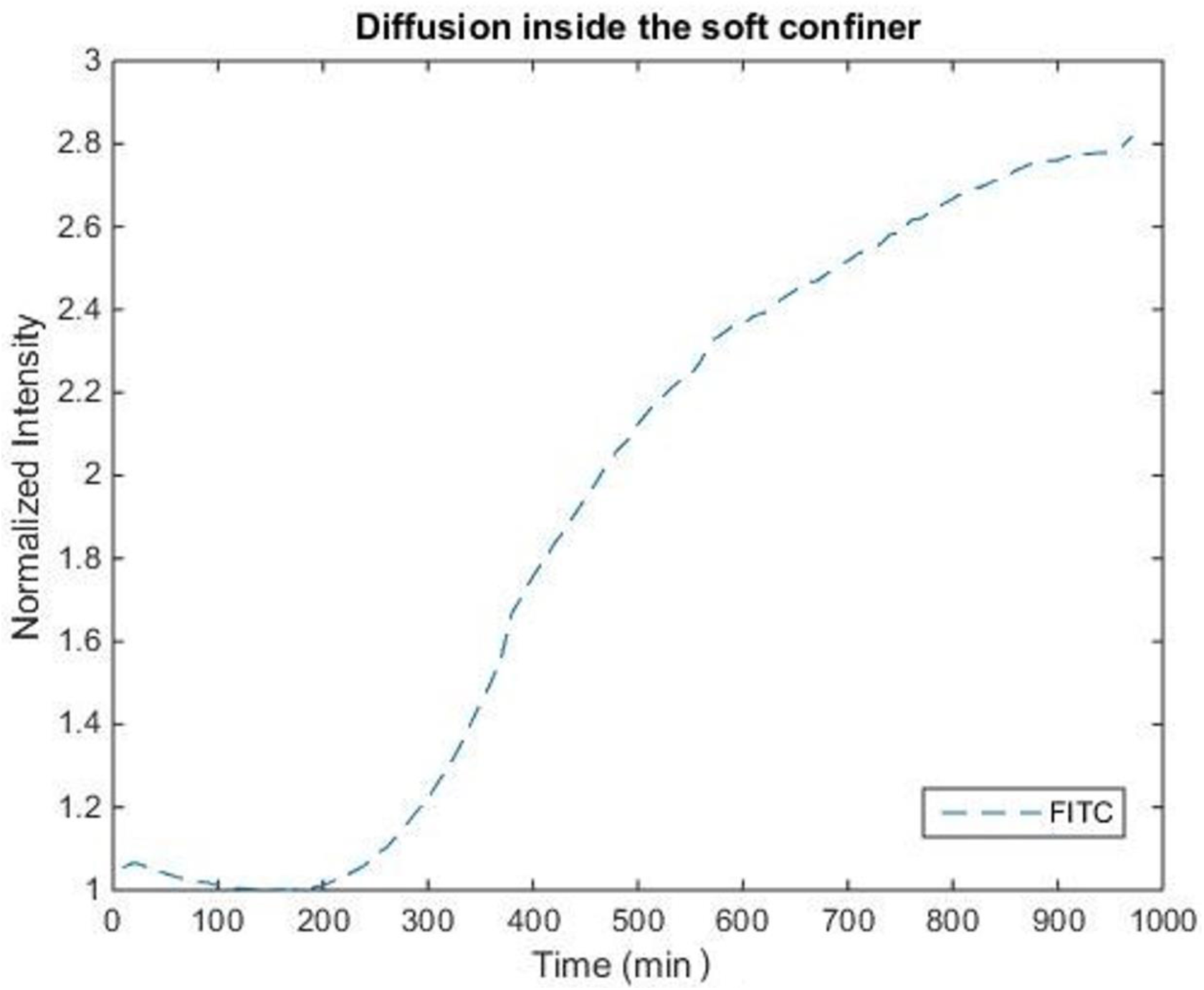
Medium diffusion inside the soft cell confiner for FITC (fluorescein isothiocyanate dissolved in PBS). Characteristic time-course of diffusion = 450 min (7 h 30 min). Briefly, the soft cell confiner was mechanically sealed and loaded with PBS. A solution of fluoresceince Isothiocyanate (0.05 μM) dissolved in PBS was then added on the open upper part of the microsystem. The fluorescence intensity in the lower part of the system was monitored every 10 min using an inverted microscope. The intensity was then measured over time and normalized against the maximum intensity.

**Figure SI 4:**
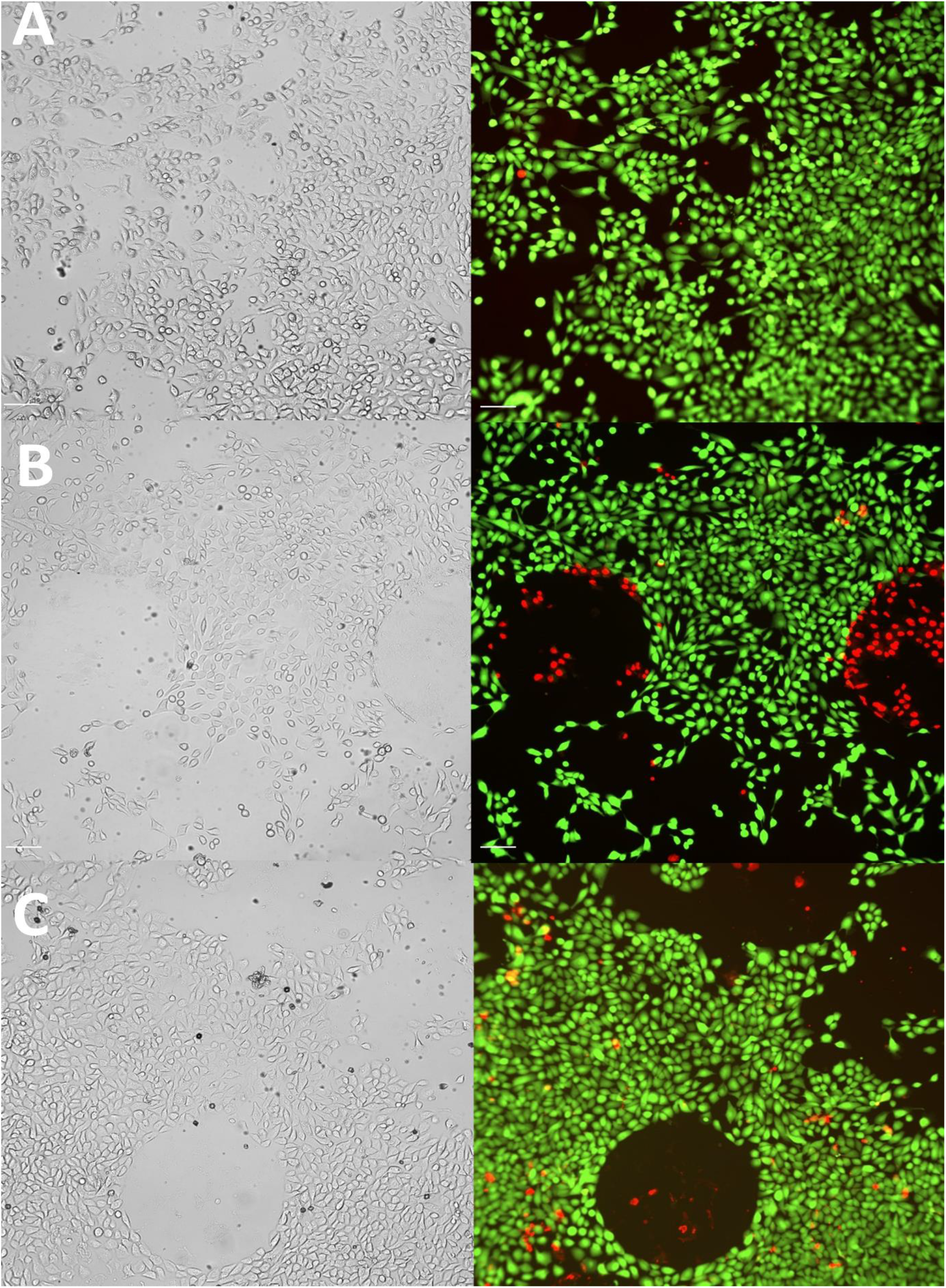
Staining of live and dead MCF10A cells after 1 day incubation in the soft cell confiner under (A) control (B) 30-μm (C) 5-μm conditions. Cells were stained for 1 h with calcein AM in green and propidium iodide in red prior to imaging. Scale bar = 100 μm.

**Movie 1:** Assembly of the soft cell confiner. Step 1: the polycarbonate holder with agarose mold installation, step 2: polycarbonate clamping washer positioning, step 3: clamping washer tightening, step 4: adding medium.

**Movie 2:** Time-lapse of phase-contrast images of immature TF1-BA hematopoietic cells over 1 day in the soft cell confiner with agarose gel presenting pillars much larger than their height (30 μm) on the left and under confinement (5 μm) on the right. Interval between images = 2 h, scale bar = 200 μm.

**Movie 3:** Time-lapse of phase-contrast images of MCF10A epithelial cells over 1 day in the confiner with agarose gel presenting pillars much larger than their height (30 μm) on the left and under confinement (5 μm) on the right. Interval between images = 1 h, scale bar = 200 μm.

## References

1 V. Vogel and M. Sheetz, Nat. Rev. Mol. Cell Biol., 2006, 7, 265–75.

2 E. K. Paluch, C. M. Nelson, N. Biais, B. Fabry, J. Moeller, B. L. Pruitt, C. Wollnik, G. Kudryasheva, F. Rehfeldt and W. Federle, BMC Biol., 2015, 13, 47.

3 E. Farge, Curr. Biol., 2003, 13, 1365–77.

4 A. J. Engler, S. Sen, H. L. Sweeney and D. E. Discher, Cell, 2006, 126, 677–89.

5 J. Du, Y. Fan, Z. Guo, Y. Wang, X. Zheng, C. Huang, B. Liang, L. Gao, Y. Cao, Y. Chen, X. Zhang, L. Li, L. Xu, C. Wu, D. A. Weitz and X. Feng, Cell Syst., 2019, 9, 214–220.e5.

6 M. Segel, B. Neumann, M. F. E. Hill, I. P. Weber, C. Viscomi, C. Zhao, A. Young, C. C. Agley, A. J. Thompson, G. A. Gonzalez, A. Sharma, S. Holmqvist, D. H. Rowitch, K. Franze, R. J. M. Franklin and K. J. Chalut, Nature, 2019, 573, 130–134.

7 D. T. Butcher, T. Alliston and V. M. Weaver, Nat. Rev. Cancer, 2009, 9, 108–22.

8 D. Wirtz, K. Konstantopoulos and P. C. P. P. C. Searson, Nat. Rev. Cancer, 2011, 11, 522.

9 M. J. Paszek, N. Zahir, K. R. Johnson, J. N. Lakins, G. I. Rozenberg, A. Gefen, C. A. Reinhart-King, S. S. Margulies, M. Dembo, D. Boettiger, D. A. Hammer and V. M. Weaver, Cancer Cell, 2005, 8, 241–254.

10 C. P. Ng, B. Hinz and M. A. Swartz, J. Cell Sci., 2005, 118, 4731–4739.

11 T. Stylianopoulos, J. D. Martin, M. Snuderl, F. Mpekris, S. R. Jain and R. K. Jain, Cancer Res., 2013, 73, 3833–3841.

12 G. Cheng, J. Tse, R. K. Jain and L. L. Munn, PLoS One, 2009, 4, e4632.

13 K. Wolf, M. Te Lindert, M. Krause, S. Alexander, J. Te Riet, A. L. Willis, R. M. Hoffman, C. G. Figdor, S. J. Weiss and P. Friedl, J. Cell Biol., 2013, 201, 1069–84.

14 M. E. Fernández-Sánchez, S. Barbier, J. Whitehead, G. Béalle, A. Michel, H. Latorre-Ossa, C. Rey, L. Fouassier, A. Claperon, L. Brullé, E. Girard, N. Servant, T. Rio-Frio, H. Marie, S. Lesieur, C. Housset, J.-L. Gennisson, M. Tanter, C. Ménager, S. Fre, S. Robine and E. Farge, Nature, 2015, 523, 92–95.

15 L. Chin, Y. Xia, D. E. Discher and P. A. Janmey, Curr. Opin. Chem. Eng., 2016, 11, 77–84.

16 S. Pradhan and J. H. Slater, Biomaterials, 2019, 215, 119177.

17 M. P. Lutolf, P. M. Gilbert and H. M. Blau, Nature, 2009, 462, 433–441.

18 E. Enzo, S. Dupont, M. Cordenonsi, F. Zanconato, S. Piccolo, M. Forcato, L. Morsut, N. Elvassore, M. Aragona, S. Bicciato, J. Le Digabel and S. Giulitti, Nature, 2011, 474, 179–183.

19 W. J. Hadden, J. L. Young, A. W. Holle, M. L. McFetridge, D. Y. Kim, P. Wijesinghe, H. Taylor-Weiner, J. H. Wen, A. R. Lee, K. Bieback, B. N. Vo, D. D. Sampson, B. F. Kennedy, J. P. Spatz, A. J. Engler and Y. S. Cho, Proc. Natl. Acad. Sci. U. S. A., 2017, 114, 5647–5652.

20 M. Kalli and T. Stylianopoulos, Front. Oncol., 2018, 8, 55.

21 L. A. Lautscham, C. Kämmerer, J. R. Lange, T. Kolb, C. Mark, A. Schilling, P. L. Strissel, R. Strick, C. Gluth, A. C. Rowat, C. Metzner and B. Fabry, Biophys. J., 2015, 109, 900–913.

22 P. Vargas, P. Maiuri, M. Bretou, P. J. Sáez, P. Pierobon, M. Maurin, M. Chabaud, D. Lankar, D. Obino, E. Terriac, M. Raab, H.-R. Thiam, T. Brocker, S. M. Kitchen-Goosen, A. S. Alberts, P. Sunareni, S. Xia, R. Li, R. Voituriez, M. Piel and A.-M. Lennon-Duménil, Nat. Cell Biol., 2015, 18, 43–53.

23 T. Lämmermann and R. N. Germain, Semin. Immunopathol., 2014, 36, 227–251.

24 J. M. Tse, G. Cheng, J. A. Tyrrell, S. A. Wilcox-Adelman, Y. Boucher, R. K. Jain and L. L. Munn, Proc Natl Acad Sci U S A, 2012, 109, 911–916.

25 D. T. Burnette, L. Shao, C. Ott, A. M. Pasapera, R. S. Fischer, M. A. Baird, C. Der Loughian, H. Delanoe-Ayari, M. J. Paszek, M. W. Davidson, E. Betzig and J. Lippincott-Schwartz, J. Cell Biol., 2014, 205, 83–96.

26 H. Kittur, W. Weaver and D. Di Carlo, Biomed. Microdevices, 2014, 16, 439–47.

27 H.-R. Thiam, P. Vargas, N. Carpi, C. L. Crespo, M. Raab, E. Terriac, M. C. King, J. Jacobelli, A. S. Alberts, T. Stradal, A.-M. Lennon-Dumenil and M. Piel, Nat. Commun., 2016, 7, 10997.

28 M. Le Berre, E. Zlotek-Zlotkiewicz, D. Bonazzi, F. Lautenschlaeger and M. Piel, Methods Cell Biol., 2014, 121, 213–229.

29 Y. J. Liu, M. Le Berre, F. Lautenschlaeger, P. Maiuri, A. Callan-Jones, M. Heuzé, T. Takaki, R. Voituriez and M. Piel, Cell, 2015, 160, 659–672.

30 V. Ruprecht, S. Wieser, A. Callan-Jones, M. Smutny, H. Morita, K. Sako, V. Barone, M. Ritsch-Marte, M. Sixt, R. Voituriez and C.-P. Heisenberg, Cell, 2015, 160, 673–85.

31 M. D. Welch, Cell, 2015, 160, 581–2.

32 A. L. McGregor, C.-R. Hsia and J. Lammerding, Curr. Opin. Cell Biol., 2016, 40, 32–40.

33 M. Raab, M. Gentili, H. de Belly, H. R. Thiam, P. Vargas, A. J. Jimenez, F. Lautenschlaeger, R. Voituriez, A. M. Lennon-Duménil, N. Manel and M. Piel, Science, 2016, 352, 359–362.

34 X. Zhang, L. Li and C. Luo, Lab Chip, 2016, 16, 1757–1776.

35 S. Zhao, Y. Chen, B. P. Partlow, A. S. Golding, P. Tseng, J. Coburn, M. B. Applegate, J. E. Moreau, F. G. Omenetto and D. L. Kaplan, Biomaterials, 2016, 93, 60–70.

36 N. W. Choi, M. Cabodi, B. Held, J. P. Gleghorn, L. J. Bonassar and A. D. Stroock, Nat. Mater., 2007, 6, 908–915.

37 A. Pathak and S. Kumar, Proc. Natl. Acad. Sci. U. S. A., 2012, 109, 10334–9.

38 M. P. Cuchiara, A. C. B. Allen, T. M. Chen, J. S. Miller and J. L. West, Biomaterials, 2010, 31, 5491–5497.

39 P. Zarrintaj, S. Manouchehri, Z. Ahmadi, M. R. Saeb, A. M. Urbanska, D. L. Kaplan and M. Mozafari, Carbohydr. Polym., 2018, 187, 66–84.

40 A. Pluen, P. a Netti, R. K. Jain and D. a Berk, Biophys. J., 1999, 77, 542–552.

41 U. Haessler, Y. Kalinin, M. a Swartz and M. Wu, Biomed. Microdevices, 2009, 11, 827–35.

42 Y. Ling, J. Rubin, Y. Deng, C. Huang, U. Demirci, J. M. Karp and A. Khademhosseini, Lab Chip, 2007, 7, 756–62.

43 S. Cosson and M. P. Lutolf, Sci. Rep., 2015, 4, 4462.

44 B. Laperrousaz, S. Jeanpierre, K. Sagorny, T. Voeltzel, S. Ramas, B. Kaniewski, M. Ffrench, S. Salesse, F. E. Nicolini and V. Maguer-Satta, Blood, 2013, 122, 3767–3777.

45 T. Kitamura, T. Tange, T. Terasawa, S. Chiba, T. Kuwaki, K. Miyagawa, Y. F. Piao, K. Miyazono, A. Urabe and F. Takaku, J. Cell. Physiol., 1989, 140, 323–34.

46 J. Goedhart, D. von Stetten, M. Noirclerc-Savoye, M. Lelimousin, L. Joosen, M. A. Hink, L. van Weeren, T. W. J. Gadella and A. Royant, Nat. Commun., 2012, 3, 751.

47 Z. N. Demou, Ann. Biomed. Eng., 2010, 38, 3509–3520.

48 Q. Chen, D. Yang, H. Zong, L. Zhu, L. Wang, X. Wang, X. Zhu, X. Song and J. Wang, Oncogenesis, 2017, 6, e375.

49 D. Gvaramia, E. Müller, K. Müller, P. Atallah, M. Tsurkan, U. Freudenberg, M. Bornhäuser and C. Werner, Biomaterials, 2017, 138, 108–117.

50 O. Lancaster, M. LeBerre, A. Dimitracopoulos, D. Bonazzi, E. Zlotek-Zlotkiewicz, R. Picone, T. Duke, M. Piel and B. Baum, Dev. Cell, 2013, 25, 270–283.

51 H. T. K. Tse, W. M. Weaver and D. Carlo, PLoS One, 2012, 7, 1–8.

52 C. J. Cattin, M. Düggelin, D. Martinez-Martin, C. Gerber, D. J. Müller and M. P. Stewart, Proc. Natl. Acad. Sci. U. S. A., 2015, 112, 11258–11263.

53 A. Elosegui-Artola, I. Andreu, A. E. M. Beedle, A. Lezamiz, M. Uroz, A. J. Kosmalska, R. Oria, J. Z. Kechagia, P. Rico-Lastres, A.-L. Le Roux, C. M. Shanahan, X. Trepat, D. Navajas, S. Garcia-Manyes and P. Roca-Cusachs, Cell, 2017, 171, 1397–1410.e14.

54 J. H. Lee, D. H. Kim, H. H. Lee and H. W. Kim, Biomaterials, 2019, 197, 60–71.

55 M. Krause, J. Te Riet and K. Wolf, Phys. Biol., 2013, 10, 065002.

56 B. Laperrousaz, L. Berguiga, F. E. Nicolini, C. Martinez-Torres, A. Arneodo, V. M. Satta and F. Argoul, Phys. Biol., 2016, 13, 03LT01.

57 T. M. Koch, S. Münster, N. Bonakdar, J. P. Butler and B. Fabry, PLoS One, 2012, 7, e33476.

58 F. J. Byfield, R. K. Reen, T.-P. Shentu, I. Levitan and K. J. Gooch, J. Biomech., 2009, 42, 1114–9.

59 H. Delanoë-Ayari, S. Iwaya, Y. T. Maeda, J. Inose, C. Rivière, M. Sano and J. P. Rieu, Cell Motil. Cytoskeleton, 2008, 65, 314–331.

60 H. Delanoë-Ayari, J. P. Rieu and M. Sano, Phys. Rev. Lett., 2010, 105, 2–5.

61 J.-P. Rieu, T. Saito, H. Delanoë-Ayari, Y. Sawada and R. R. Kay, Cell Motil. Cytoskeleton, 2009, 66, 1073–86.

62 J.-P. Rieu and H. Delanoë-Ayari, Phys. Biol., 2012, 9, 066001.

63 T. a Ulrich, A. Jain, K. Tanner, J. L. MacKay and S. Kumar, Biomaterials, 2010, 31, 1875–84.

64 G. C. Ingavle, S. H. Gehrke and M. S. Detamore, Biomaterials, 2014, 35, 3558–3570.

65 Y. P. Singh, N. Bhardwaj and B. B. Mandal, ACS Appl. Mater. Interfaces, 2016, 8, 21236–21249.

66 S. H. Medina, B. Bush, M. Cam, E. Sevcik, F. W. DelRio, K. Nandy and J. P. Schneider, Biomaterials, 2019, 202, 1–11.

67 A. Marturano-Kruik, A. Villasante, K. Yaeger, S. R. Ambati, A. Chramiec, M. T. Raimondi and G. Vunjak-Novakovic, Biomaterials, 2018, 150, 150–161.

68 A. Nicolas-Boluda, A. K. A. Silva, S. Fournel and F. Gazeau, Biomaterials, 2018, 150, 87–99.

